# Rationally designed modular STAT-activating scaffolds enforce cell-intrinsic transcriptional programs augmenting the anti-tumor potency of CAR T cells

**DOI:** 10.1101/2023.12.22.573170

**Authors:** Christopher P Saxby, Taylor K Ishida, James M Rosser, Benjamin C Curtis, Michael L Baldwin, Cailyn H Spurrell, Justin Giles, Ardizon Valdez, Joshua Gustafson, Adam J Johnson, Jacob S Appelbaum, Michael C Jensen

## Abstract

Chimeric antigen receptor (CAR)-expressing T cells can mediate anti-tumor responses in a variety of preclinical models and clinical settings, however, strategies to enhance anti-tumor potency is the subject of intense investigation. Signals emanating from gamma-c cytokine receptors modulate the transcriptional state of activated T cells impacting proliferation, survival, differentiation, and effector functioning through the STAT family of transcription factors. Design of ligand-independent cell-intrinsic cytokine STAT activation scaffolds is a conceptually attractive strategy to provide CAR T cells with a surrogate for exogenous cytokine support. Here, we designed a series of ligand-autonomous STAT inducer (LASI) scaffolds comprised of an extracellular identification tag, a homodimerizing transmembrane domain, and a membrane proximal IL7R Box1 domain followed by STAT5 and/or STAT3 docking sequences derived from IL7R and IL21R, respectively. We constructed LASI scaffolds having STAT5 (LASI-5), STAT3 (LASI-3), and combined STAT5 and STAT3 (LASI-5+3) docking domains and then interrogated their impact in primary human CD8^+^ anti-CD19 (4-1BB:zeta) CAR T cells. While LASI-5 expression had limited effects on CAR T cells, LASI-3 transcriptional programming was found to be indispensable to achieving anti-tumor functional enhancement associated with limited terminal differentiation, heightened T cell proliferation in response to antigen, and dampened expression of exhaustion-associated genes. Moreover, CAR T cells supplemented with LASI-3 or 5+3 displayed superior potency against human leukemia tumors in NSG mice. LASI-5+3 mediated the highest magnitude of CAR T cell engraftment *in vivo* that evolved into a fatal lymphoproliferative syndrome. However, the same efficacy enhancement was achieved with LASI-3 without the lymphoproliferative complication. Our findings provide a rationale for utilization of constitutively expressed LASI-3 to enhance the anti-tumor potency of CAR T cells, the need to regulate the activity of LASI-5+3, and a generalizable scaffold design for studying additional combinations of STAT family transcription factors.

## Introduction

Cytokines are small, secreted proteins that mediate immunomodulation through autocrine, paracrine, and endocrine signaling. During an immune response, cytokines allow myriad cell types to share information through patterns of cytokine secretion, receptor expression and downstream signal transduction. Lymphocytes rely on the gamma chain family of cytokines for development and modulation, and the eponymous gamma chain receptor is required for signaling by all members of this cytokine family. Furthermore, gamma chain cytokines play a pivotal role in the T cell response, as T cell activation and co-stimulation requires a third signal of inflammatory cytokines to elicit acquisition of effector function, survival, and memory formation.^1^ In immuno-oncology, T cells can be programmed using CARs for activation and costimulation in response to target antigen engagement. However, without direct control of cytokine signaling, the CAR T cell response depends on the peri-tumoral cytokine milieu, which often lacks pro-inflammatory cytokines. Next generation CAR therapies will require synthetic biomedical engineering strategies to provision T cells with cytokine signals that support engraftment, persistence, and durable tumor regressions.

Gamma chain cytokine signaling are mediated, in part, by JAK-STAT signaling and downstream transcriptional networks. Signal transduction occurs when an extracellular cytokine induces the association of the common gamma chain with one or more specific cytokine receptors. Box 1 and Box 2 motifs in the intracellular domains of these cytokine receptors bind JAK proteins, which, when in proximity due to cytokine-induced receptor oligomerization, transphosphorylate one another.^2^ Thus activated JAKs phosphorylate key tyrosine residues in the cytokine cytoplasmic domains, which in turn become docking sites for STAT proteins. Docked STATs become phosphorylated by JAK proteins, leading to STAT dimerization and translocation to the nucleus, where they become active transcription factors. There are seven mammalian STAT proteins, each with a unique DNA binding specificity and subsequent pattern of transcriptional activity.

Gamma chain cytokines mediate distinct effects on T cell function and differentiation by driving unique patterns of STAT activation. IL-2 and IL-7, for example, primarily activate STAT5, which is responsible for promoting T cell homeostatic proliferation^3^ and antigen activation-driven clonal expansion.^4^ Another member of the gamma chain cytokine family, IL-21 promotes effector functioning and negatively regulates T cell exhaustion in chronic viral infection models,^5,6^ an effect mediated by IL-21-induced activation of STAT3.^7^ STAT5 and STAT3 therefore each offer distinct effects on T cell biology.

Favorable outcomes from CAR T cell therapy are correlated with greater T cell expansion and persistence^8^ and lower expression of markers associated with T cell exhaustion.^9^ We hypothesized that orthogonal activation of STAT5 and STAT3 would impart CAR T cells with greater persistence and functional durability, respectively. To promote these desirable T cell characteristics, we constructed a ligand-autonomous STAT5 and STAT3 inducer (LASI-5+3). We generated this scaffold protein by modifying a constitutive STAT-5-activating IL7R mutant^10^ with a peptide from the IL21R responsible for activating STAT3.^11^

To parse the effects STAT3 and STAT5 signaling on CAR T cell functionality, we iterated on the LASI-5+3 design to develop a panel of constructs with variable levels of STAT3 and STAT5 activation. Subsequent *in vitro* assays demonstrated that STAT-activating scaffolds imparted improved CAR T cell proliferation, survival, and anti-tumor durability. Furthermore, STAT3 activation preserved a subset of T cells with a stem cell memory phenotype, downregulated expression of exhaustion-associated genes, and promoted expression of quiescence-associated genes. In a xenograft model of B cell leukemia, only STAT3-supplemented CAR T cells demonstrated complete tumor clearance and marked improvements in mouse survival. Here, we present a hybrid STAT-signaling technology with marked improvements to CAR T cell potency. LASI implementation provides insights into the effects of specific cytokine signals on CAR T cell therapeutic functionality and offers a promising platform for tailoring cytokine signaling profiles to fit the therapeutic niches of specific indications in the future.

## Materials and Methods

### Construct Design

The ligand-autonomous STAT inducer (LASI) sequences included four variants: Marker-Only control, LASI-5, LASI-5+3, and LASI-3. The Marker-Only control sequence consisted of a truncated version of Her2, Her2tG.^12^ The LASI-5 sequence featured the Her2tG extracellular domain, a mutant IL7R transmembrane domain with a cysteine-containing insertion,^10^ and the native IL7R intracellular domain. LASI-5+3 was constructed by appending a peptide sequence from the IL21R to the end of the LASI-5 intracellular domain sequence via a flexible glycine-serine linker, (G_4_S)_2_. Finally, the LASI-3 variant was created by mutating the LASI-5+3 sequence to replace the tyrosine sequence at position 449 in the native IL7R sequence with a sequence encoding phenylalanine. All LASI sequences were introduced into a construct featuring a piggyBac transposon for integration into the genome.^13^ The piggyBac construct harbored two divergent coding sequences: 1) A polycistronic sequence driven by the human elongation factor 1 alpha (EF1a) promoter encoding the human G01S anti-CD19CAR,^14^ followed by a P2A ribosomal skip sequence,^15^ a double mutant of the dihydrofolate reductase (DHFRdm) for drug selection,^16^ a T2A ribosomal skip sequence, and finally a truncated epidermal growth factor receptor (EGFRt) to serve as cell surface marker;^17^ and 2) an upstream sequence encoding each LASI variant and driven by the MND promoter.^18^ Sequences were procured and codon-optimized using the Invitrogen GeneArt gene synthesis service, and fragments were assembled into the piggyBac plasmid backbone using Gibson assembly.^19^

### T Cell Production and Culture

Protocols to acquire human cells were approved by the institutional review board of Seattle Children’s Hospital. CD4^+^ and CD8^+^ T cells were isolated from whole blood using a STEMCELL RoboSep-S automated cell separation instrument according to manufacturer’s protocols. Flow-through from the T cell sorting was subjected to a LymphoPrep (STEMCELL, Cat. # 07811) overlay centrifuge-based protocol for the isolation of residual peripheral blood mononucleated cells (PBMCs). PiggyBac transposon constructs and RNA encoding transposase were introduced into T cells by electroporation using a Lonza 4D-Nucleovector instrument and the P3 Primary Cell Kit (Lonza, Cat. # V4XP-3032). Electroporated T cells were immediately seeded into 24-well G-Rex plates (Wilson Wolf, Cat. # 80240M) along with donor-matched PBMC feeder cells. Culture media consisted of X-Vivo 15 (Lonza, Cat. # BP04-744Q) supplemented with 2% KnockOut Serum Replacement (ThermoFisher, Cat. # 10828-028) by volume, 4.6ng/mL IL-2 (STEMCELL, Cat. # 78220.3), 20ng/mL IL-4 (Miltenyi, Cat. # 130-093-924), 10ng/mL IL-7 (Miltenyi, Cat. # 130-095-363) and 20ng/mL IL-21 (Miltenyi, Cat. # 130-095-784). Unmodified T cell control conditions were cultured in the absence PBMCs and treated with a TransAct (Miltenyi, Cat. # 130-111-116) CD3/CD28 stimulation. Three days later, drug selection of modified T cells was begun by adding 50nM methotrexate (Accord, Cat. # NDC 16729-277-30) to the culture medium. Twenty-one days after culture initiation, T cells were collected and cryopreserved in CryoStor CS5 (STEMCELL, Cat. # 07933) for subsequent studies. All post-production assays were conducted in complete RPMI media (cRPMI), which consisted of RPMI 1640 (Gibco, Cat. # 22400-089) supplemented with 10% FBS (Hyclone, Cat. # SH30071.03) and 2mM L-glutamine (Gibco, 25030-081). Unless otherwise noted, all subsequent studies were conducted after mixing CD4^+^ and CD8^+^ T cell groups to achieve a 1:1 starting ratio.

### Flow Cytometry

Flow cytometric analysis was performed to determine transduction marker expression and phenotypic surface marker expression using the following generalized protocol. Cells were removed from culture and placed into 96-well round-bottom plates, washed twice with PBS (Gibco, Cat. # 10010-023), stained with antibodies specific for cell surface markers, washed two more times with PBS and finally fixed with 0.5% paraformaldehyde (Electron Microscopy Sciences, Cat. # 15713) in PBS before analysis. Flow cytometric data was collected using a BD LSRFortessa flow cytometer and later analyzed using FlowJo software. Final flow plots were populated by cell populations remaining after the following gating strategy was performed. First, a “lymphocyte” gate was generated by drawing a polygon within the forward scatter vs side scatter scatterplot to isolate events with size and granularity characteristic of lymphocytes and remove debris and dead cell events. Within the lymphocyte gate, a second “single cell” gate was generated by gating on events that followed a linear relationship between forward scatter in the height and area dimensions to remove cell doublets from downstream analysis. If applicable, a “live cells” gate was generated using the live dead stain to exclude cells with compromised membranes. Specific cell staining reagents are listed below, organized by context of the assay.

The transgenic surface marker panel during T cell production consisted of: Erbitux APC (BD, Cat. # 624367), Herceptin PE (BD Cat. # 624255), Anti-CD3 BUV395 (BD, Cat. # 563546), Anti-CD4 BV785 (BioLegend, Cat. # 317442), Anti-CD8 Pacific Blue (BioLegend, Cat. # 344718) Live Dead Fixable Aqua viability dye (Thermo, Cat # L34957).

The weekly tumor challenge panel for T cell tracking consisted of: Anti-CD4 APC-Cy7 (BioLegend, Cat. # 317418), Anti-CD8 Pacific Blue (BioLegend, Cat. # 344718), Fixable Viability Stain 520 (BD, 564407).

The T cell phenotypic characterization panel during tumor challenge assays consisted of: Anti-CD4 BV785 (BioLegend, Cat. # 317442), Anti-CD8 Pacific Blue (BioLegend, Cat. # 344718) Live Dead Fixable Aqua viability dye (Thermo, Cat # L34957), Anti-CCR7 FITC (BioLegend, Cat. # 353215), Anti-CD45RA BV786 (BD, Cat. # 741010), Anti-CD45RO APC (BioLegend, Cat. # 304210), Anti-CD62L PE (BioLegend, Cat. # 304806).

The T cell activation and exhaustion marker panel during tumor challenge assays: Anti-CD4 BV785 (BioLegend, Cat. # 317442), Anti-CD8 Pacific Blue (BioLegend, Cat. # 344718) Live Dead Fixable Aqua viability dye (Thermo, Cat # L34957), Anti-TIGIT APC (BioLegend, Cat. # 372705).

### Intracellular Cytokine Staining

T cells were co-cultured with mCherry-expressing Raji tumor cells at a 2:1 effector to tumor ratio for 6 hours. Immediately after cells were plated, a CD107a-specific flow cytometry antibody (BioLegend, Cat. # 328640) was added to the culture to detect degranulation. Two hours after the co-culture began, a transport inhibitor cocktail (ThermoFisher, Cat. # 00-4980-03) was added to prevent T cell release of secretory proteins. At the end of co-culture, the cells were incubated with 10 µL of Fc block (Miltenyi, Cat. # 130-059-901) for 10 minutes at room temperature to avoid indiscriminate antibody binding by tumor cells. Cells were then stained with Live Dead Fixable Near IR viability dye (ThermoFisher, Cat # L10119), anti-CD4 BUV737 (BD, Cat # 612748) and anti-CD8 Pacific Blue (BioLegend, Cat. # 344718) to allow gating of live T cells. Next, samples were fixed and permeabilized using the Fixation/Permeabilization Kit (BD, Cat. # 554714) according to the manufacturer’s protocol. After permeabilization, the cells were stained with antibodies specific for secretory proteins: Granzyme B (BD, Cat. # 396407), Interferon-gamma (BD, Cat. # 563731), TNF-alpha (BD, Cat. # 563996), IL-2 (BioLegend, Cat. # 500326), and Perforin (BioLegend, Cat. # 353303). Flow cytometric data was collected using a BD LSRFortessa flow cytometer and analyzed using FlowJo software.

### Phospho-Flow Cytometry

CD4^+^ T cells were harvested from culture fourteen days after electroporation and seeding with PBMC feeder cells. The cells were washed to remove residual cytokine from the culture medium and replated in cytokine-free media for 48 hours. After cytokine-free rest, the cells were moved into FACS tubes and stained with Live Dead Fixable Aqua Viability Dye (Invitrogen, Cat. # L34966). Cells were then resuspended in pre-warmed cRPMI media, and control groups received media with to 5ng/mL IL-7 (Miltenyi, Cat. # 130-095-363) or 5ng/mL IL-21 (Miltenyi, Cat. # 130-095-784) added. After a 20-minute incubation at 37° C, the cells were fixed using pre-warmed CytoFix (BD, Cat. # 554655), permeabilized using pre-chilled Perm Buffer III (BD, Cat. # 558050) and stained with anti-phosphoSTAT3 Alexa Fluor 647 (BD, Cat. # 557815) or anti-phosphoSTAT5 Alexa Fluor 647 (BD, Cat. # 612599). Samples were analyzed within 16 hours of staining to avoid signal loss.

### Repeated CAR T Cell Stimulation Assays

The following protocol describes how T cells were subjected to recursive tumor stimulations, either on a weekly or a three-day basis. Twenty-one days after initial electroporation, CD4^+^ and CD8^+^ CAR T cells were mixed at a 1:1 ratio and seeded for repeated tumor challenge assays. On day 0, mCherry-expressing Raji tumor cells were added to T cell cultures at a 2:1 effector to target ratio. Exogenous cytokine control cultures were supplemented with 5ng/mL IL-7 (Miltenyi, Cat. # 130-095-363), 5ng/mL IL-21 (Miltenyi, Cat. # 130-095-784), or both. At either day 3 or day 7, samples of each co-culture were harvested, and live cell counts were taken using a Cellaca MX cell counting instrument (Nexcelom Biosciences). Flow cytometry was performed, as described in the “flow cytometry” section above, to assess the composition of the culture between CD4^+^ T cells, CD8^+^ T cells, and mCherry^+^ Raji tumor cells. Live T cell counts were back-calculated using the percent live cells in the culture accounted for by T cells multiplied by the total live cells according to cell counts. T cells were then reseeded in fresh media and tumor cells were again added at a 2:1 effector to target ratio. The counting and flow analysis process was repeated one additional time at the end of the assay. T cell growth curves were assembled by calculating fold expansion of each T cell group across both tumor stimulations.

### Incucyte Cytotoxicity Assays

CAR T cell cytotoxicity was evaluated over a variable period of days using the S3 IncuCyte Live Cell Imager (Sartorius). To begin, ten thousand mCherry-expressing Raji tumor cells were cultured per well of a flat-bottom 96-well plate. Next, T cells were added at effector to tumor ratios of 2:1, 1:1, 0.5:1 or 0.25:1. Unless otherwise specified, all groups were cultured in cytokine-free media for these studies. Exogenous cytokine control cultures were supplemented with 5ng/mL IL-7 (Miltenyi, Cat. # 130-095-363), 5ng/mL IL-21 (Miltenyi, Cat. # 130-095-784), or both. Fluorescent and phase images of these co-cultures were collected every four hours. IncuCyte image analysis software was used to quantify tumor presence over time by mCherry signal. Simultaneously, Incucyte “Cell by Cell” analysis software was used to discriminate “low-red” objects for tracking of T cell presence over the course of the assay. Two experimental schemas were employed: 1) T cell groups received one tumor challenge at day 0 and imaging proceeded for six days, and 2) T cell groups received five tumor challenges every three days and imaging proceeded for fifteen days. In the multi-challenge schema, ten thousand tumor cells were added per well during each challenge.

### Cytokine-Free Growth and Apoptotic Status Assay

To assess cytokine-free T cell growth in the absence of antigen stimulation, T cell groups were subjected to a single tumor exposure and monitored for growth thereafter. CAR T cells were co-cultured with irradiated mCherry-expressing Raji cells at a 1:1 effector to target ratio. Seven days later, cultures were washed to remove secreted cytokines, flow cytometry confirmed the absence of residual mCherry^+^ tumor cells, and T cells were returned to culture. Ten days after initial tumor exposure, T cells were harvested to assess cell health and apoptotic status using the PE Annexin V Apoptosis Detection Kit I (BD, Cat. # 559763) according to manufacturer’s instructions. Live T cell concentrations were monitored intermittently thereafter until day 23 using a Cellaca MX cell counting instrument (Nexcelom Biosciences).

### RNA-seq

LASI-expressing CD19CAR T cells were co-cultured with mCherry-expressing Raji tumor cells at a 2:1 effector to target ratio in cRPMI media. One week later, the cells were removed from culture and labeled with anti-CD4 APC-Cy7 (BioLegend, Cat. # 317418) and anti-CD8 Pacific Blue (BioLegend, Cat. # 344718) antibodies and a viability dye (BD, Cat. # 564407). Labeled cell populations were sorted for live CD4^+^ and CD8^+^ T cells using a Sony MA900 cell sorter, excluding mCherry^+^ Raji tumor cells. Sorted cell populations were snap-frozen in liquid nitrogen and stored at -80° C.

RNA was isolated from cell pellets using Directzol RNA Miniprep (Zymo) including the optional DNase treatment. RNA quality was assessed using a 4150 TapeStation (Agilent) with an RNA 6,000 Nano Kit (Agilent). RNA sequencing libraries were made using the TruSeq Stranded mRNA kit (Illumina) according to manufacturer instructions. The resulting RNA-seq libraries’ quality and concentration were assessed using the D1000 DNA Kit (Agilent) on the TapeStation and a Qubit Fluorometer with the Qubit dsDNA HS Assay Kit (Life Technologies). RNA-seq libraries were pooled and sequenced via paired end 150 bp reads using Genewiz’s sequencing only service (HiSeq or Novaseq 6000) at a target depth of 25M reads per library (Illumina). RNA-seq was attempted on all cytokine samples, with libraries for 37 samples passing minimum quality metrics for inclusion in the analysis. RNA-seq data was analyzed using the nf-core RNA-seq pipeline version 3.4.^20^ Briefly, reads were mapped to the human genome (GRCh38) using STAR aligner (v2.6.1d). Gene expression counts were generated using salmon (v1.5.2).

### Differential gene expression

Differential gene expression analysis between two phenotypes (LASI vs cytokine) was performed using edgeR version 3.32.1.^21^ First, genes whose expression was very low in most samples were discarded from further analyses. Next, *estimateCommonDisp* and *estimateTagwiseDisp* were run to properly handle over-dispersion at the global and single gene level. Across samples, normalization was then performed using TMM, via the function *calcNormFactors*. Differentially expressed genes were determined using the *exactTest* function, as those showing the multiple testing corrected FDR <0.05, a two-sided raw p value <0.01 and |log_2_(fold change)| >=1. The total number of expressed genes includes all genes present in at least two samples with normalized CPM (counts per million) of at least 1. PCA was performed using the log2-transformed, TMM-normalized expression values as input. PCA results were robust to the choice of the most variable genes used as input (range tested: 1,000 to 10,000).

### Leukemia Xenograft Mouse Studies

T cells were cryopreserved for *in vivo* studies after being mixed to achieve a ∼3:2 ratio of CD4^+^ to CD8^+^ T cells. 11-13 week old NOD scid gamma (NSG) immunodeficient mice were injected via the tail vein with one million human leukemia Nalm-6 tumor cells, modified to express a fusion protein of mCherry and firefly luciferase. Six days after tumor injection, tumor engraftment in each mouse was quantified via bioluminescent imaging using an IVIS Spectrum In Vivo Imaging System (Perkin Elmer) and Living Image Software. Mice were then distributed into treatment groups to normalize average engraftment across groups as much as possible. Seven days after tumor injection, mice were injected via the tail vain with four million T cells. Thereafter, mice were monitored for tumor growth by bioluminescent imaging. Health metrics and weight changes were also recorded to capture effects of different T cell treatment groups. Mice were euthanized when they showed moderate to severe hind-limb paralysis, a result of unchecked tumor progression, or otherwise as recommended by veterinary staff. Beginning on day 17 and at weekly intervals thereafter, retro-orbital bleeds were performed to examine T cell engraftment in the peripheral blood.

### T Cell Quantification in Retro-Orbital Bleed Samples

Peripheral blood samples were taken by retro-orbital bleeds of mice on a weekly basis, and the following flow cytometry protocol was performed to quantify T cells in the blood. 40uL of blood was moved into a 96 well plate. Remaining blood was pooled across samples and mixed with T cells from culture to serve as an FMO control staining mix. Other controls included T cell only samples and Nalm-6 only samples to allow for discrimination of these populations in the blood during analysis. Red blood cells were lysed using Pharm Lyse Buffer (BD, Cat. # 555899), and samples were treated with Fc blocking reagent (Miltenyi, Cat. # 130-059-901) to prevent indiscriminate antibody binding. Next, cells were stained with the following panel of reagents: anti-CD3 BUV737 (BD, Cat. # 612751), anti-human CD45 APC-Cy7 (BD, Cat. # 557833), anti-mouse CD45 PerCP/Cy5.5 (BioLegend, Cat. # 103132), and fixable viability stain 520 (BD, Cat. # 564407). Finally, samples were fixed in 0.5% paraformaldehyde (Electron Microscopy Sciences, Cat. # 15713) in PBS before analysis. CountBright absolute counting beads (Invitrogen, Cat. # 2207530) were added to each sample to allow for back-calculation of T cell counts per µL of blood analyzed. Flow cytometric data was collected using a BD LSRFortessa flow cytometer and analyzed using FlowJo software.

## Results

### Modularly designed STAT-activating scaffolds provide constitutive STAT5 and STAT3 signaling in CAR T cells

We developed a hybrid cytokine signaling technology in the form of a ligand-autonomous STAT inducer protein for constitutive STAT5 and STAT3 activation in T cells (LASI-5+3). The LASI-5+3 protein is composed of four key domains, each imparting a distinct functionality (**Fig. 1a**). The extracellular portion of LASI-5+3 features a Her2 domain recognized by the Herceptin antibody, allowing for direct detection of the scaffold protein on the cell surface.^12^ The transmembrane domain is composed of an IL7R transmembrane domain variant harboring a cysteine insertion, which facilitates auto-homodimerization via a disulfide bridge leading to constitutive signaling.^10^ Intracellularly, LASI-5+3 features the full-length IL7R intracellular domain and a tethered IL21R peptide to provide STAT5 and STAT3 signaling, respectively.

**Figure 1:**
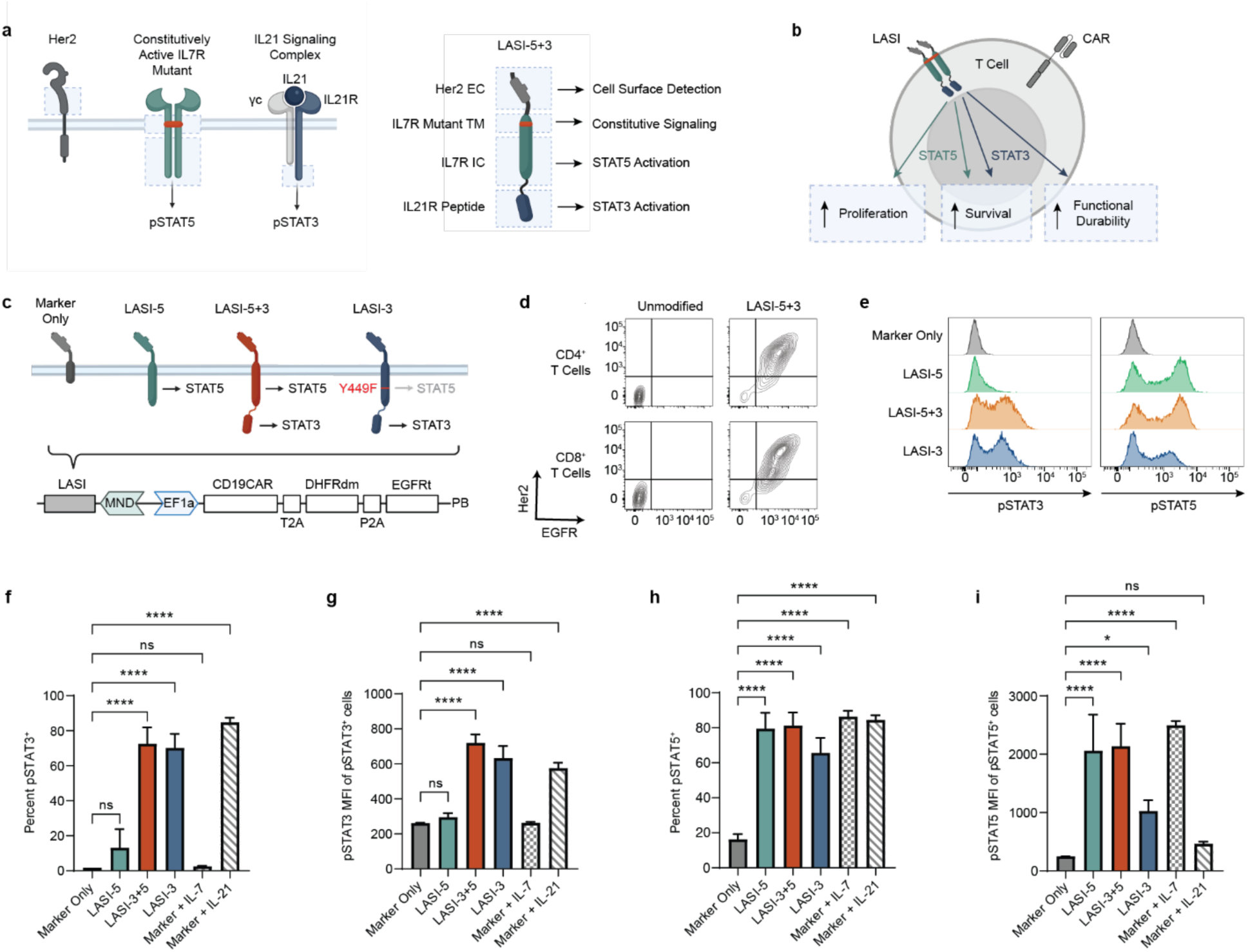
Modularly designed STAT-activating scaffolds provide constitutive STAT5 and STAT3 signaling in CAR T cells. **a**, Construction of a ligand-autonomous STAT inducer (LASI-5+3) from elements of Her2, IL7R and IL21R for constitutive activation of STAT3 and STAT5. **b**, Hypothetical effects of LASI technology on CAR T cell expansion, persistence, and anti-tumor function. **c**, Panel of LASI variants designed for specific STAT signaling outputs, and introduction of these variants into a divergent dual-promoter construct for co-expression with the anti-CD19CAR in T cells. **d**, Flow cytometry analysis of surface expression of LASI, marked by Her2tG, and the anti-CD19CAR polycistron, marked by EGFRt. **e**, Flow cytometry analysis of phosphorylated STAT3 and STAT5 (pSTAT3 and pSTAT5) in LASI-expressing CAR T cells. Histograms display cell distributions from a representative donor. **f**, Percent pSTAT3^+^ cells across three donors. **g**, Average pSTAT3 median fluorescent intensity (MFI) among pSTAT3^+^ cells across three donors. **h**, Percent pSTAT5^+^ cells across three donors. **i**, Average pSTAT5 MFI among pSTAT3^+^ cells across three donors. *Statistics: n = 3 biological replicates. Plots show mean +/- standard deviation. One-way ANOVA was performed with multiple comparisons to Marker-Only group. p < 0.05* 0.01** 0.001*** 0.0001\*\*\*\**.

STAT5 and STAT3 activation both promote T cell survival,^22–24^ but activation of STAT5 specifically supports T cell proliferation,^3^ whereas activation of STAT3 specifically confers durable effector function.^7^ We therefore hypothesized that supplementing CAR T cells with LASI expression would bolster CAR T cell therapeutic potency by: 1) supporting robust CAR T cell expansion, 2) enhancing CAR T cell survival and persistence, and 3) promoting durable anti-tumor activity (**Fig. 1b**). To parse the effects STAT3 and STAT5 activation on CAR T cell functionality, we leveraged the modular design of LASI-5+3 to develop a panel of constructs with different STAT signaling outputs (**Fig. 1c**). First, we included a “Marker-Only” construct featuring no intracellular signaling domains as a control. Next, we modified LASI-5+3 to generate two further variants: 1) Removing the IL21R peptide yielded LASI-5 for diminished STAT3 activation capacity, and 2) ablating a STAT5 activation motif in the IL7R intracellular domain yielded LASI-3 for diminished STAT5 activity.

These four LASI variants were inserted into a divergent dual-promoter construct for simultaneous expression with a human anti-CD19 CAR.^14^ Introduction of these constructs into CD4^+^ and CD8^+^ T cells resulted in dual-positive populations expressing LASIs, as marked by Her2tG, and the anti-CD19CAR-encoding polycistron, as marked by EGFRt (**Fig. 1d, Extended Data Fig. 1a-c**). Next, we examined the ability of LASI-expressing cells to activate STATs via flow cytometry to detect the phosphorylated versions of STAT3 and STAT5 (pSTAT3 and pSTAT5) (**Fig. 1e**). As expected, compared to Marker-Only control, only LASI-5+3 and LASI-3 demonstrated significant activation of STAT3, both in terms of percent pSTAT3^+^ cells (**Fig. 1f**) and in the pSTAT3 MFI of those pSTAT3^+^ cells (**Fig. 1g**). Patterns of STAT3 activation by LASI7/21 and LASI-3 were comparable to treatment of Marker-Only cells with exogenous IL-21.

STAT5 activation patterns proved to be more complex, with all LASI variants and cytokine treatments resulting in a significant increase in percent pSTAT5^+^ compared to Marker-Only (**Fig. 1h**). Examination of pSTAT5 MFI among pSTAT5^+^ cells revealed that LASI-5 and LASI-5+3 provided the highest magnitude of STAT5 activation, approximately equal to levels in Marker-Only cells treated with exogenous IL-7 (**Fig. 1i**). Unexpectedly, despite the mutation of the tyrosine responsible for STAT5 signaling in IL7R, LASI-3 retained a reduced but significant level of STAT5 activation, suggesting an alternative means of STAT5 activation. Having developed a panel of STAT inducer proteins with three distinct signaling patterns, we went on to investigate how each variant impacted CAR T cell functionality.

### STAT3-activating scaffolds promote quiescence-associated gene expression and sustain a subset of stem cell memory CAR T cells

To investigate potential gene expression differences promoted by LASI supplementation, we performed RNA sequencing on CD8^+^ CAR T cells. We prepared T cells for sequencing by stimulating with CD19-expressing Raji tumor cells followed by sorting CD8^+^ T cell populations one week later (**Fig. 4a**). A list of differentially expressed genes (DEGs) was assembled by comparing normalized transcript levels between each LASI group and the Marker-Only control (**Fig. 4b**). As expected, given its common signaling outputs, LASI-5+3- expressing T cells showed the greatest number of DEGs in common with the other two LASI groups. However, LASI-3-expressing CAR T cells showed the greatest number of DEGs, suggesting that STAT3 activation with low-level STAT5 activation leads to the greatest changes in gene expression. Furthermore, genes downregulated in LASI-3-expressing T cells compared to the Marker-Only group included exhaustion-associated genes TOX2, NR4A2 and NR4A3^25^ (**Fig. 4c**). LASI-3 supplementation was also marked by increased expression of KLF2 and KLF3, transcription factors promoting T cell quiescence.^26,27^

Given this modulation of transcription factors involved in T cell exhaustion and activation, we examined the effect of LASIs on the expression of key differentiation markers of naïve T cell populations *in vitro*.^28^ To model T cell differentiation in response to antigen exposure, we repetitively stimulated CAR T cell groups with Raji tumor cells every three days and tracked expression of differentiation markers CD45RA, CD62L and CCR7 by flow cytometry (**Fig. 4d**). We found that STAT3-signaling LASIs, and particularly LASI-3, showed higher percentages of CD8^+^ T cells expressing markers of stem cell memory T cells (**Fig. 4e**). The differences in marker expression between STAT3-signaling LASIs and other T cell groups widened between the first and second tumor exposures, suggesting that the effects of LASI-5+3 and LASI-3 on differentiation became more apparent as CAR T cells are repetitively activated. In addition, treatment of Marker-Only cells with IL-21 had a similar effect of preserving a population of T cells with stem cell memory phenotype.

To examine potential effects of LASIs on T cell exhaustion markers, we concurrently examined TIGIT expression as CAR T cells were repeatedly exposed to antigen. TIGIT is an inhibitory receptor marking exhausted T cells^29^ and associated with CAR T cell dysfunction in treatment of non-Hodgkin lymphoma.^30^ While no statistically significant differences emerged, we observed a trend that STAT3-signaling LASIs as well as treatment with IL-21 led to decreased TIGIT expression (**Extended Data Fig. 2a**), suggesting that equipping CAR T cells with STAT3-signaling LASIs may decrease susceptibility to T cell exhaustion and deactivation via TIGIT ligation. Taken as a whole, our findings demonstrated that expression of LASI-3 and, to a lesser degree, LASI-5+3 maintained a quiescent, stem cell memory phenotype and decreased expression of genes involved in T cell exhaustion.

### STAT inducers increase CAR T cell proliferation and suppress apoptosis

Proliferative capacity and longevity are key features of potent CAR T cell products, as an effective anti-tumor response requires that CAR T cells expand *in situ* upon encountering tumor cells and persist until tumor can be eradicated. We examined the effects of LASI expression on CAR T cell proliferation and survival *in vitro* after co-culture with antigen-bearing tumor cells. When subjected to weekly exposures to Raji cells (**Fig. 3a**), LASI-5+3- and LASI-3-expressing CAR T cells displayed significantly enhanced proliferation (**Fig. 3b, c**). In keeping with this finding, Marker-Only cells cultured with IL-21 alone or IL-7 and IL-21 likewise showed a proliferative benefit. Unexpectedly, LASI-3- expressing CAR T cells demonstrated greater expansion magnitudes than all other groups, including Marker-Only cells cultured with exogenous IL-7 and/or IL-21. Augmented proliferation in LASI-5+3 and LASI-3-expressing CAR T cells did not result in skewing toward CD4^+^ of CD8^+^ populations, as these groups roughly maintained the starting 1:1 ratio of CD4^+^ to CD8^+^ T cells after two weeks in culture (**Extended Data Fig. 2b**). Therefore, LASIs with STAT3 signaling activity provide enhanced CD4^+^ and CD8^+^ T cell proliferation *in vitro*, roughly mimicking the effects of treatment with exogenous cytokines. While Marker-Only T cells required both IL-7 and IL-21 supplementation to achieve maximal expansion, the greatest proliferative boost was conferred by LASI-3, suggesting that diminished STAT5 signaling may be optimal for augmented antigen-dependent proliferation. Furthermore, the pro-proliferative effects appeared to stem directly from LASI signaling rather than downstream cytokine secretion, as endogenous IL-2, TNF-⍺ and IFN-*γ* levels were not significantly affected by LASI expression (**Extended Data Fig. 2c, d**).

**Figure 2:**
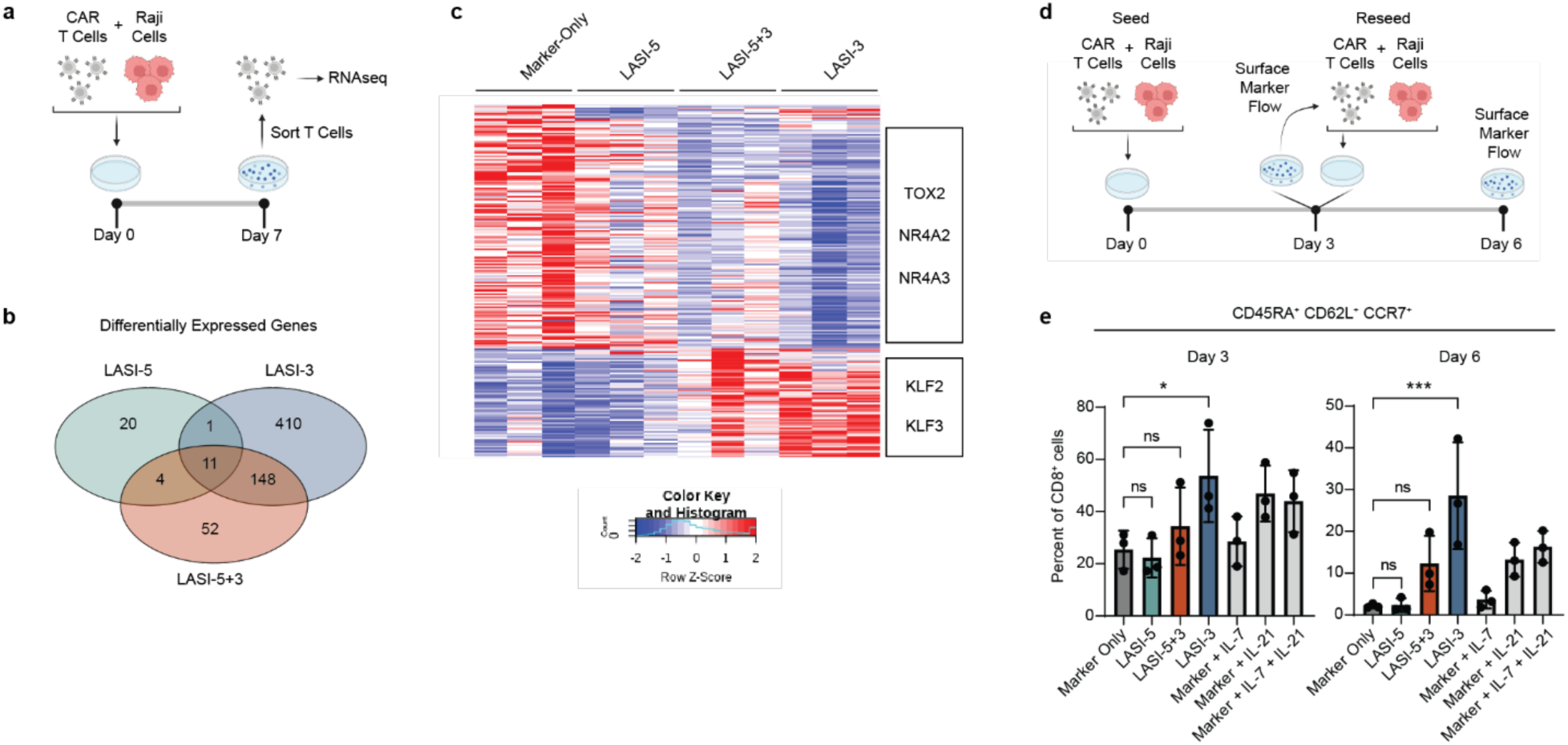
STAT3-activating scaffolds promote quiescence-associated gene expression and sustain a subset of stem cell memory CAR T Cells. **a**, Experimental schema to prepare CAR T cells for RNA-seq. **b**, differentially expressed genes (DEGs) in CD8^+^ T cells across LASI groups compared to Marker-Only control. DEG list includes all genes with a multiple-testing corrected FDR < 0.05, a two-sided raw p-value < 0.01 and log_2_(fold change)| >=1. **c**, Heatmap displaying DEGs arrayed by expression pattern across T cell groups with designated positions of genes of interest. **d**, Experimental approach of to track T cell phenotypic markers upon two stimulations with Raji tumors every three days. **e**, Percent of CD8^+^ T cell populations expressing CD45RA, CD62L and CCR7 at day 3 and day 6. *Statistics: n = 3 biological replicates. Plots show mean +/- standard deviation. One-way ANOVA was performed with multiple comparisons to Marker-Only group. p < 0.05* 0.01** 0.001\*\*\**.

**Figure 3:**
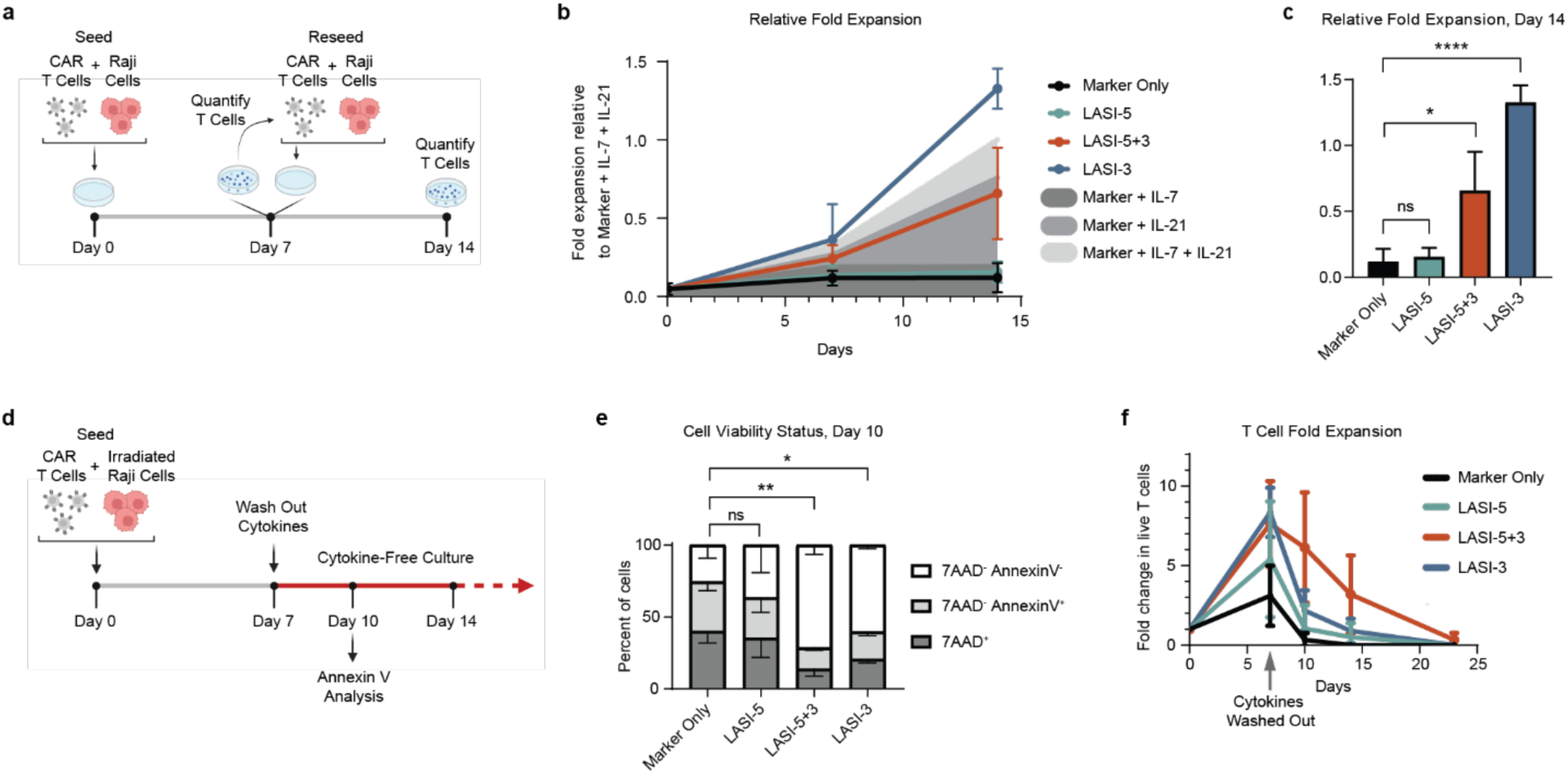
STAT inducers increase CAR T cell proliferation and suppress apoptosis. **a**, Experimental schema to assess LASI-supplemented CAR T cell growth upon two exposures to Raji tumor cells one week apart. **b**, Average relative fold expansion of T cells across three donors. All fold expansion values were normalized on a donor-by-donor basis to the Marker Only + IL-7 + IL-21 condition to account for differences in baseline proliferative capacity across donors. The shaded areas indicate the average relative expansion of Marker Only cells treated with cytokines. **c**, Average relative fold expansion at day 14. **d**, Experimental schema to assess T cell growth and survival in absence of cytokines and CAR antigen. **e**, Cell viability status ten days after stimulation with irradiated Raji tumors, assessed by flow cytometry to detect cell incorporation of 7-AAD and Annexin V. **f**, Fold change in live T cells after stimulation with irradiated Raji cells. *Statistics: n = 3 biological replicates. Plots show mean +/- standard deviation. One-way ANOVA was performed with multiple comparisons to Marker-Only group. p < 0.05* 0.01** 0.001*** 0.0001\*\*\*\**.

To examine T cell proliferation and survival in the absence of cytokine *and* antigen, we subjected T cell groups to a single stimulation with irradiated Raji cells, followed by a wash-out of secreted cytokines seven days later, at which point all Raji tumor cells had been eliminated (**Fig. 3d**). Assessing cell viability status three days after cytokine wash-out revealed that LASI-5+3- and LASI-3-expressing T cells showed significantly decreased proportions of apoptotic and pre-apoptotic cells, identified as 7-AAD^+^ and 7-AAD^-^/Annexin-V^+^ populations, respectively (**Fig. 3e**). This pro-survival effect did not correlate with BCL2 expression, as revealed by RNA-seq in CD8^+^ T cells (**Extended Data Fig. 2e**). However, LASI-5+3 and LASI-3-expressing cells showed significantly lower expression of the pro-apoptotic gene BCL2L11, a potential mechanism for maintaining T cell viability. T cells from all groups exhibited complete attrition of viable cells over the course of several weeks in the absence of cytokines and antigen, with LASI-5+3 providing the greatest extension in T cell survival (**Fig. 3f**). As with proliferation, STAT3 signaling distinguishes LASIs that exhibit a significant pro-survival effect *in vitro* in the absence of cytokine and CAR antigen. However, the superiority of LASI-5+3 in extending T cell persistence suggests that pairing STAT3 and STAT5 activation may yield the greatest resistance to apoptosis.

### STAT3-signaling scaffolds improve CAR T cell anti-tumor activity under prolonged tumor exposure

In addition to proliferation and survival, we investigated the ability of LASIs to augment CAR T cell anti-tumor activity. We used live cell fluorescent imaging to track mCherry-marked Raji tumor cells in co-culture with CAR T cells at a range of effector to target (E:T) cell ratios. In an assay of short-term cytotoxicity, we co-cultured T cells and Raji cells and monitored fluorescence for six days thereafter (**Extended Data Fig. 3a**). Comparing the tumor signal in each CAR T cell group to that of unmodified T cells allowed us to calculate the percent tumor suppression over the six-day period (**Extended Data Fig. 3b**). While LASI-5+3 and LASI-3 outperformed other groups at most E:T ratios, only LASI-3 conferred a statistically significant enhancement in tumor suppression at the 0.5:1 E:T (**Extended Data Fig. 3c**). Expression of cytotoxic proteins GZMB and PRF as well as degranulation marker CD107A in CD8^+^ T cells did not vary significantly across LASI groups (**Extended Data Fig. 3d**), suggesting an alternative mechanism for the enhanced anti-tumor function provided by LASI-3. Treatment of Marker-Only CAR T cells with cytokines yielded similar changes in tumor suppression to LASI supplementation. However, in contrast to the dominance of LASI-3 amongst the LASI-bearing T cell groups, treatment of Marker-Only cells with IL-21 alone did not lead to greater tumor suppression than treatment with IL-7 and IL-21. As with proliferative capacity, the superior anti-tumor performance of LASI-3 suggested that diminished STAT5 signaling may be optimal for short-term anti-tumor activity afforded by LASIs.

Looking to investigate effects of LASI expression on T cell cytotoxicity on a longer timescale, we exposed T cell groups to series of five “tumor challenges” every three days (**Fig. 4a**). Longitudinal tracking of tumor signal revealed that STAT3-signaling LASIs conferred a significant enhancement in tumor suppression by CAR T cells (**Fig. 4b, c**), and that these trends were mimicked by treatment of Marker Only control cells with exogenous cytokines (**Fig. 4d**). Furthermore, simultaneous tracking of T cell populations showed a correlated increase in T cell expansion by STAT3-signaling LASIs over the fifteen-day period (**Fig. 4e, f, g**). Taken together, these findings suggest that STAT3 activation, either by LASI expression or treatment with exogenous IL-21, leads to durable anti-tumor activity *in vitro*.

**Figure 4:**
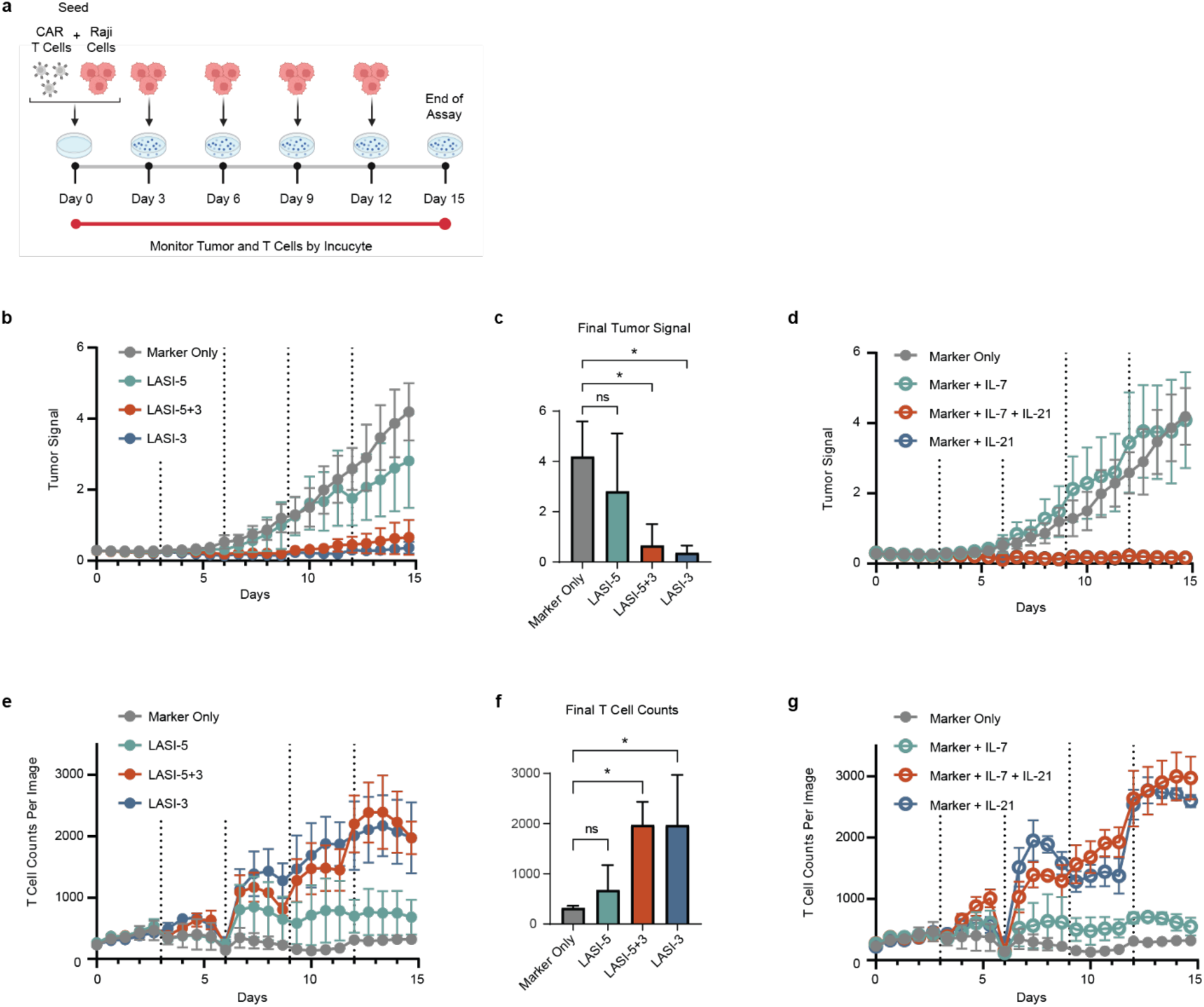
STAT3-signaling scaffolds improve CAR T cell anti-tumor activity under prolonged tumor exposure. **a**, Schema of a long-term cytotoxicity assay with five recursive tumor challenges with Raji cells every three days. **b**, Longitudinal tumor signal in LASI-supplemented T cell groups. Dashed vertical lines indicate times when fresh tumor cells were to the co-culture. **c**, Average final tumor signal across LASI-supplemented groups. **d**, Longitudinal tumor signal in Marker-Only cytokine-treated T cell groups. **e**, Longitudinal T cell counts in LASI-supplemented T cell groups. **f**, Average final T cell counts across LASI-supplemented groups. **g**, Longitudinal T cell counts in Marker-Only cytokine-treated groups. *Statistics: n = 3 biological replicates. Plots show mean +/- standard deviation. One-way ANOVA was performed with multiple comparisons to Marker-Only group. p < 0.05\**.

### STAT inducers support CAR T cell mediated elimination of leukemia tumors *in vivo*

*In vitro* studies have highlighted the impacts of LASI expression on CAR T cell proliferation, survival, anti-tumor potency, and differentiation. We went on to examine whether these features affect the ability of CAR T cells to eliminate tumors *in vivo*. To this end, we employed a human leukemia xenograft model in NSG mice using CD19^+^ Nalm-6 tumor cells modified to express firefly luciferase for bioluminescent tracking (**Fig. 5a**). CD4^+^ and CD8^+^ T cells were modified with LASI/CD19CAR constructs for *in vivo* studies (**Extended Data Fig. 4a**) and mixed to achieve ∼3:2 CD4^+^ to CD8^+^ ratio across T cell groups (**Extended Data Fig. 4b**). T cells were administered one week after tumor injection, and T cell engraftment and persistence were monitored by *ex vivo* flow cytometry on peripheral blood. As expected, mice treated with any anti-CD19CAR T cell product showed a delay in tumor progression compared to mice treated with Unmodified T cells (**Fig 5b**). However, all mice treated with LASI-5+3- or LASI-3-expressing CAR T cells showed a complete elimination of tumor signal and return background flux values. LASI-5 CAR T cells achieved greater reductions in tumor burden than Marker-Only CAR T cells but failed to achieve full tumor eradication in any mice.

**Figure 5:**
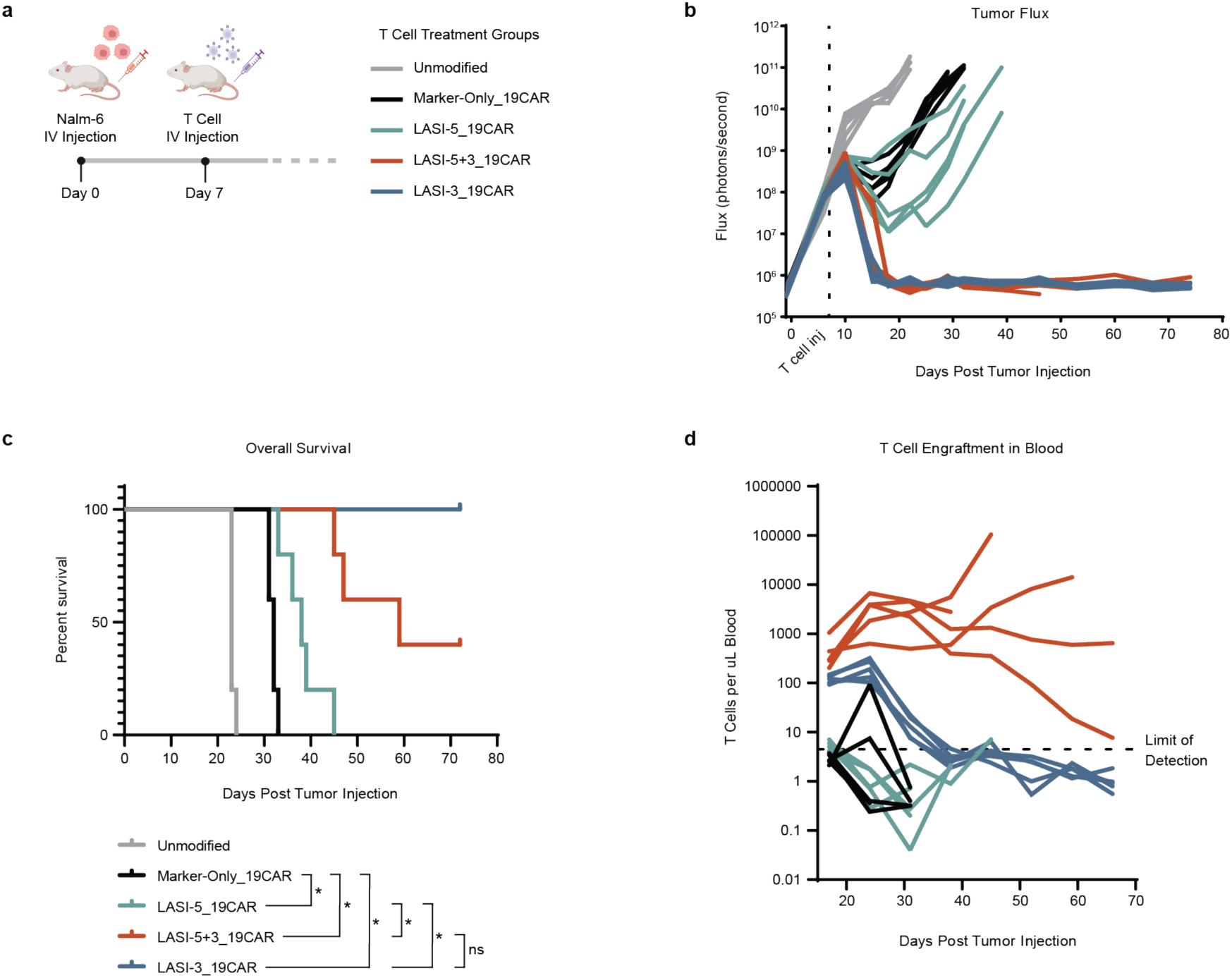
STAT inducers support CAR T cell mediated elimination of leukemia tumors *in vivo.* **a**, Experimental schema of human Nalm-6 leukemia xenograft *in vivo* model (n = 5 mice per group). **b**, Tumor tracking by bioluminescent imaging of mice. Each line represents an individual mouse. **c**, Kaplan Meier survival curve with accompanying statistics to determine significance (Mantel-Cox test, Bonferroni-corrected threshold: p < 0.0083*). **d**, Tracking of T cell concentration in peripheral blood. Each line represents an individual mouse.

These differences in tumor control led to significant effects on mouse survival (**Fig. 5c**), with LASI-5+3 and LASI-3 extending survival beyond other treatment groups. However, unexpectedly, several mice treated with LASI-5+3 CAR T cells developed abdominal swelling and became moribund despite absence of tumor signal, necessitating euthanasia. Necropsy of moribund mice revealed abdominal swelling to originate from enlarged liver and spleen suggesting a pathological role of circulating CAR T cells. LASI-5+3 CAR T cells appeared at the highest concentrations in peripheral blood early in the study and remained elevated throughout (**Fig. 5d**), especially in mice suffering from liver and spleen enlargement. Given that LASI-5+3 CAR T cells continued to expand and persist in mice after tumor had been eliminated, we hypothesize that moribundity arose in this group due to unrestricted T cell expansion and engraftment in the liver and spleen, eventually leading to organ damage. This phenomenon was only observed in the LASI-5+3 treatment group at a penetrance of 60%.

In contrast, while LASI-3 CAR T cells appeared at elevated concentrations compared to Marker-Only CAR T cells at early timepoints, their frequency fell below the limit of detection by day 38 post-tumor injection. We conclude that the combination of constitutive STAT3 and high STAT5 signaling in LASI-5+3 triggers unrestricted CAR T cell growth after tumor has been eliminated. High STAT5 activation appeared to be dispensable for LASI enhancement of anti-tumor responses, however, as LASI-3 CAR T cells achieved the same level of tumor control as LASI-5+3 without causing CAR T cell antigen-independent lymphoproliferation in mice.

## Discussion

The modular design of our STAT-activating scaffold allowed us to generate variants with differential STAT5 and STAT3transcriptional outputs. *In vitro* studies revealed that LASIs providing STAT3 activation led to the most pronounced improvements in T cell proliferation, survival, and functional durability. Surprisingly, LASI-5+3 did not outperform LASI-3 on these metrics, suggesting that high STAT5 activation is dispensable to mediate functional enhancements to CAR T cells. Rather, in some cases, high STAT5 activation appeared to counteract effects of STAT3 activation, as LASI-3 showed greater impacts on T cell differentiation markers and gene expression than LASI-5+3.

*In vivo*, however, high STAT5 activation by LASI-5+3 yielded more notable effects on CAR T cell behavior. Though LASI-5 supplementation did not confer significant functional enhancements *in vitro*, LASI-5-expressing CAR T cells increased tumor suppression and survival *in vivo*. In addition, LASI-5+3-equipped CAR T cells displayed the highest T cell concentrations in the peripheral blood. Perhaps the unremarkable effects of high STAT5 activation *in vitro* can be attributed to an abundance of endogenous cytokines secreted by CAR T cells, such as IL-2 and IL-15. The presence of these endogenous cytokines may have created an elevated background level of STAT5 activation, effectively diluting the effect of STAT5 activity of LASI origin.

Only LASIs providing STAT3 activation resulted in complete elimination of leukemia tumors *in vivo*, and this effect can likely be attributed to the heightened T cell engraftment achieved by LASI-5+3 and LASI-3. Other effects observed *in vitro*, such as delayed T cell differentiation and downregulation of exhaustion-related genes, may have also played a role. However, it is unlikely that differences in T cell differentiation status emerged during the T cell production culture period, as all T cell groups showed similar differentiation markers at the end of culture before the initiation of *in vivo* studies (**Extended Data Fig. 4c**).

The combination of STAT3 and high STAT5 activation provided by LASI-5+3 resulted in unrestricted T cell growth after tumors were eliminated *in vivo*. STAT3 activation appears to license T cells for robust expansion upon encounter with tumor *in vivo*, but high STAT5 activation is required for sustaining T cell populations after antigen-bearing tumor diminishes. This finding is keeping with cytokine literature, as IL-7 has been shown to promote homeostatic proliferation of T cells,^31^ an effect that can be recapitulated by the introduction of a constitutively active variant of STAT5.^3^ The utilization of LASI-5+3 would require engineering of regulated expression strategies to merit further development as a therapeutic module.

LASI-3 mediated durable anti-tumor responses without triggering unchecked T cell growth, making it the most attractive candidate to supplement CAR T cell products. Future work will examine other signaling cascades triggered by LASI-3 beyond STATs, including AKT and ERK pathways, to elucidate the networks responsible for the functional enhancements promoted by this protein. The hybrid cytokine signaling platform detailed here allows systematic investigation of combinatorial cytokine signaling, yielding insights into the interplay between STAT3 and STAT5 in CAR T cells. Our technology licenses future efforts to engineer adoptive T cell products with custom cytokine support to meet the barriers imposed by specific tumor types.

## Extended Data Figures

**Extended Data Figure 1:**
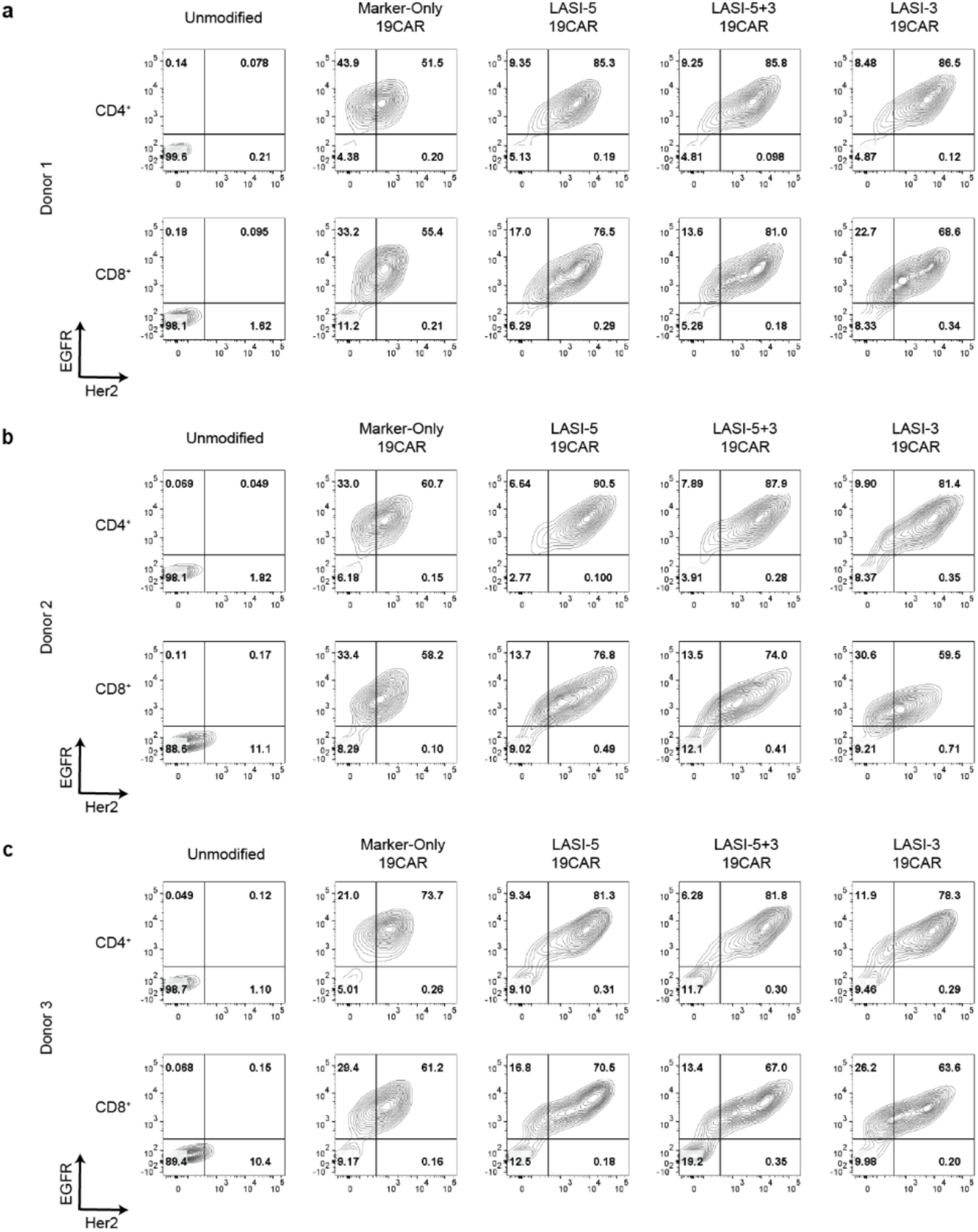
Transgenic cell surface marker expression in T cells generated from three donors for *in vitro* studies. Flow cytometry contour plots showing cell surface expression of transgenic markers in CD4^+^ and CD8^+^ cells modified with divergent dual-promoter constructs. EGFR marks expression of the anti-CD19CAR-containing polycistron, and Her2 marks expression of LASI variants. Plots display marker expression in three donors (donor 1 in **a**, donor 2 in **b**, and donor 3 in **c**) used as biological replicates for *in vitro* assays.

**Extended Data Figure 2:**
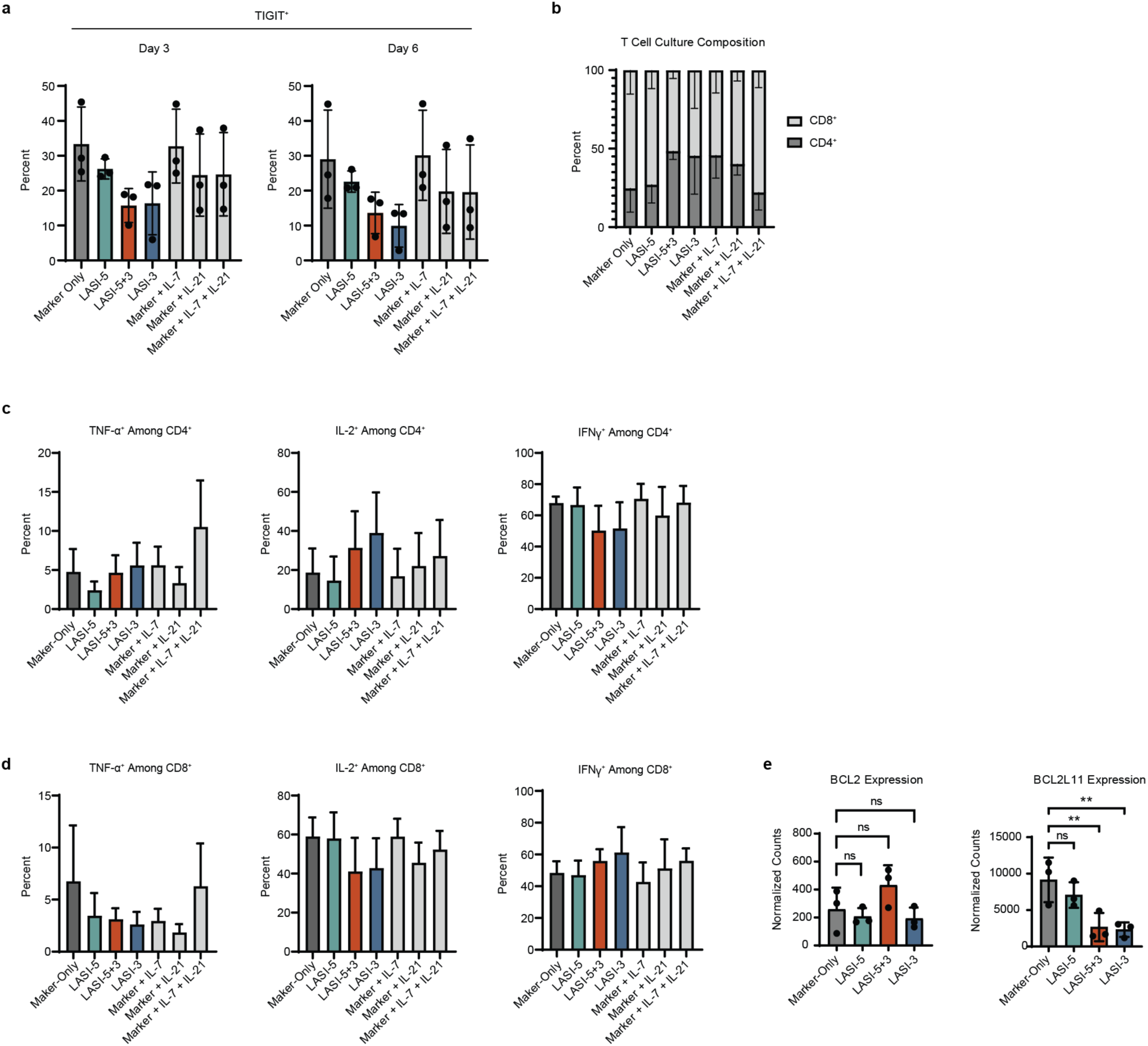
Effects of LASI expression on TIGIT expression, CD4^+^ vs. CD8^+^ culture distribution, cytokine expression and expression of apoptosis-regulating proteins in CAR T cells. **a**, Percent of CD8^+^ T cell populations expressing TIGIT at day 3 and day 6 of the experiment outlined in Figure 2d. **b**, Average percent CD4^+^ and CD8^+^ populations in T cell culture at day 14 of the experiment outlined in Figure 3a. **c**, Percent of CD4^+^ CAR T cells expressing TNF-⍺, IL-2, and IFN-*γ* when co-cultured with Raji tumor cells, as determined by flow cytometry with intracellular cytokine staining. **c**, Percent of CD8^+^ CAR T cells expressing TNF-⍺, IL-2, and IFN-*γ* when co-cultured with Raji tumor cells. **e**, Normalized transcript counts of BCL2 and BCL2L11 genes by RNA-seq experiment outlined in Figure 2a. *Statistics (sub-figures a, c, and d): n = 3 biological replicates. Plots show mean +/- standard deviation. One-way ANOVA was performed with multiple comparisons to Marker-Only group. Sub-figures a, c, and d: no statistically significant differences were found. Sub-figure e: p < 0.01\*\**.

**Extended Data Figure 3:**
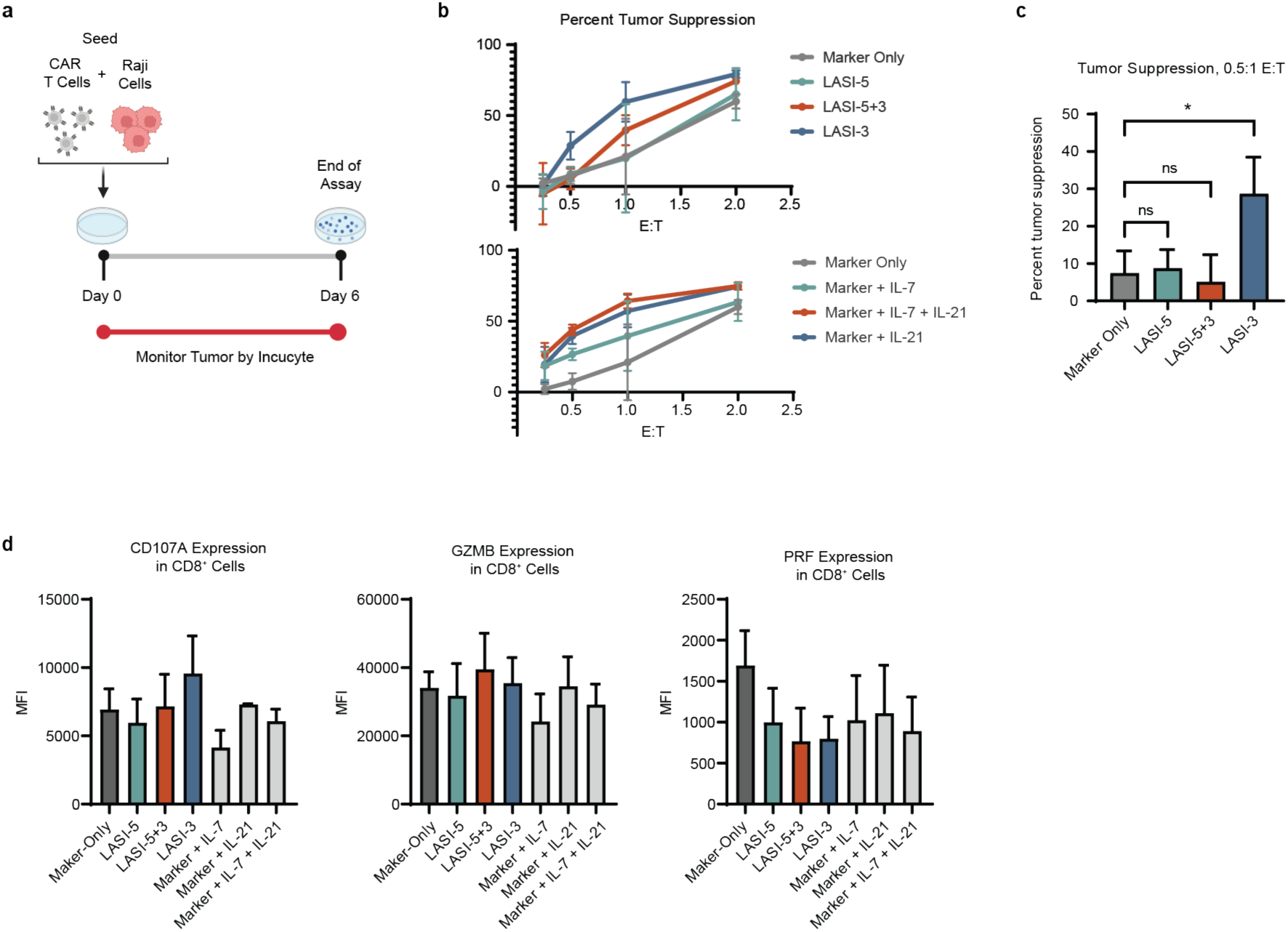
Effects of LASI expression on short-term cytotoxicity and cytotoxic protein expression. **a**, Schema of a short-term cytotoxicity assay using live cell fluorescent imaging. **b**, Percent tumor suppression at end of assay by LASI-expressing T cell groups (top) and Marker-Only T cell groups (bottom) across various effector to target (E:T) ratios. **c**, Percent tumor suppression by LASI-expressing groups at 0:5:1 E:T. **d**, Median fluorescence intensity (MFI) of cytotoxic proteins GZMB and PRF as well as degranulation marker CD107A in CD8^+^ CAR T cells as determined by flow cytometry with intracellular and cell surface staining. *Statistics: n = 3 biological replicates. Plots show mean +/- standard deviation. One-way ANOVA was performed with multiple comparisons to Marker-Only group. Sub-figure c: p < 0.05*. Sub-figure d: no statistically significant differences were found*.

**Extended Data Figure 4:**
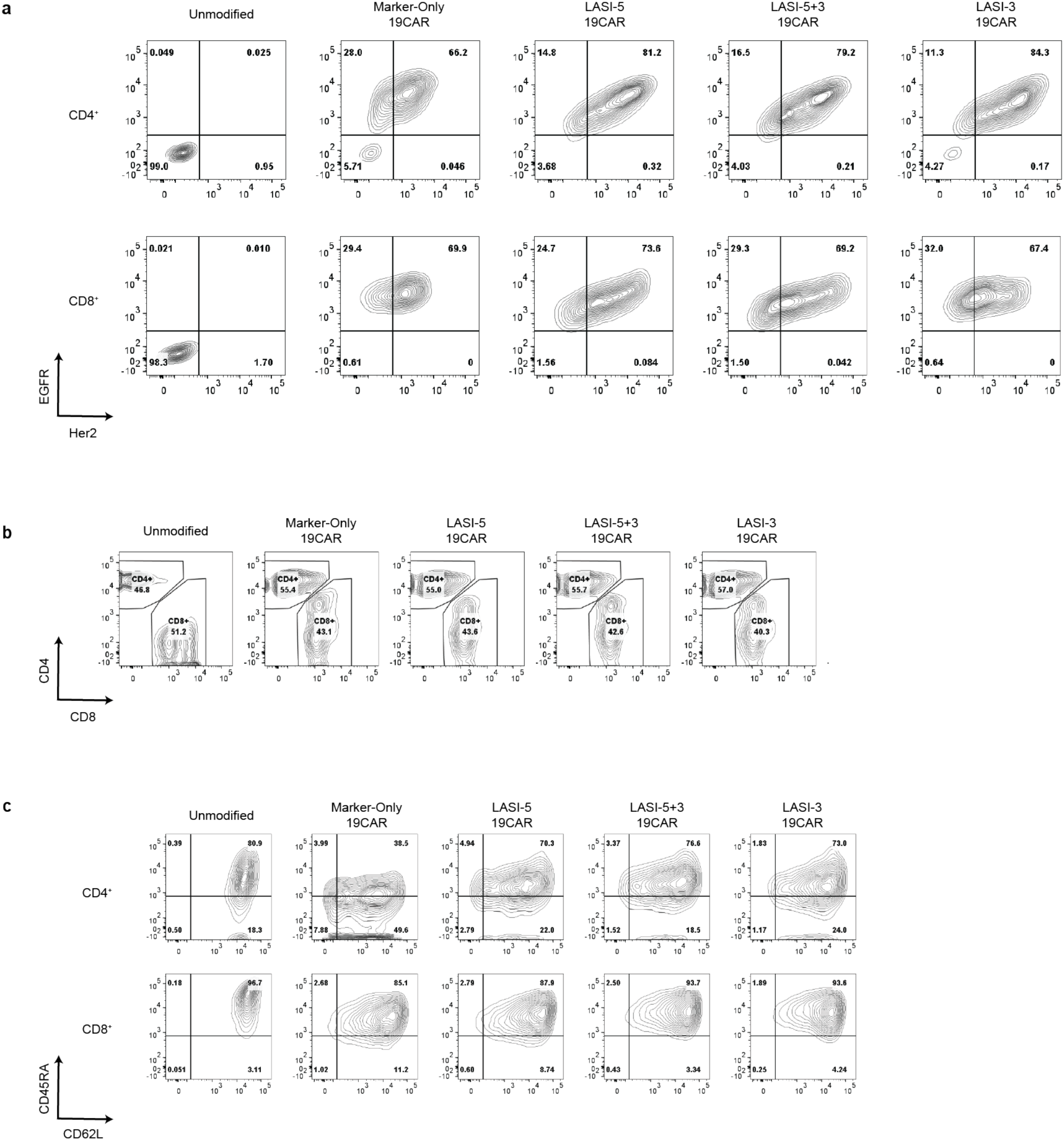
Transgenic cell surface marker and phenotypic marker expression by T cells generated for *in vivo* studies. **a**, Flow cytometry contour plots showing constitutive cell surface expression of transgenic markers in CD4^+^ and CD8^+^ cells modified with divergent dual-promoter constructs for *in vivo* studies. EGFRt marks expression of the anti-CD19CAR-containing polycistron, and Her2tG marks expression of LASI variants. **b**, CD4^+^ and CD8^+^ T cell cultures were mixed to normalize the CD4:CD8 ratio across groups before cryopreservation in preparation for *in vivo* studies. Contour plots show CD4 and CD8 expression in each T cell group after thaw for *in vivo* studies. **c**, Expression of differentiation markers CD45RA and CD62L in CD4^+^ and CD8^+^ T cells produced for *in vivo* studies.

## References

1. Curtsinger, J. M. & Mescher, M. F. Inflammatory cytokines as a third signal for T cell activation. Curr Opin Immunol 22, 333–340 (2010).

2. Morris, R., Kershaw, N. J. & Babon, J. J. The molecular details of cytokine signaling via the JAK/STAT pathway. Protein Science 27, 1984–2009 (2018).

3. Burchill, M. A. et al. Distinct Effects of STAT5 Activation on CD4+ and CD8+ T Cell Homeostasis: Development of CD4+CD25+ Regulatory T Cells versus CD8+ Memory T Cells. The Journal of Immunology 171, 5853–5864 (2003).

4. Grange, M. et al. Activated STAT5 promotes long-lived cytotoxic CD8 +T cells that induce regression of autochthonous melanoma. Cancer Res 72, 76–87 (2012).

5. Yi, J. S., Du, M. & Zajac, A. J. A Vital Role for Interleukin-21 in the Control of a Chronic Viral Infection. Science (1979) 324, 1572–1576 (2009).

6. Fröhlich, A. et al. IL-21R on T cells is critical for sustained functionality and control of chronic viral infection. Science 324, 1576–80 (2009).

7. Xin, G. et al. A Critical Role of IL-21-Induced BATF in Sustaining CD8-T-Cell-Mediated Chronic Viral Control. Cell Rep 13, 1118–1124 (2015).

8. Cappell, K. M. & Kochenderfer, J. N. Long-term outcomes following CAR T cell therapy: what we know so far. Nat Rev Clin Oncol 20, 1–13 (2023).

9. Majzner, R. G. & Mackall, C. L. Clinical lessons learned from the first leg of the CAR T cell journey. Nat Med 25, 1341–1355 (2019).

10. Shum, T. et al. Constitutive Signaling from an Engineered IL7 Receptor Promotes Durable Tumor Elimination by Tumor-Redirected T Cells. Cancer Discov 7, 1238–1247 (2017).

11. Zeng, R. et al. The molecular basis of IL-21–mediated proliferation. Blood 109, 4135–4142 (2007).

12. Johnson, A. J. et al. Rationally Designed Transgene-Encoded Cell-Surface Polypeptide Tag for Multiplexed Programming of CAR T-cell Synthetic Outputs. Cancer Immunol Res 9, 1047–1060 (2021).

13. Li, X. et al. piggyBac transposase tools for genome engineering. Proceedings of the National Academy of Sciences 110, (2013).

14. Sommermeyer, D. et al. Fully human CD19-specific chimeric antigen receptors for T-cell therapy. Leukemia 31, 2191–2199 (2017).

15. Liu, Z. et al. Systematic comparison of 2A peptides for cloning multi-genes in a polycistronic vector. Sci Rep 7, 2193 (2017).

16. Jonnalagadda, M. et al. Efficient selection of genetically modified human T cells using methotrexate-resistant human dihydrofolate reductase. Gene Ther 20, 853–60 (2013).

17. Wang, X. et al. A transgene-encoded cell surface polypeptide for selection, in vivo tracking, and ablation of engineered cells. Blood 118, 1255–1263 (2011).

18. Robbins, P. B. et al. Increased probability of expression from modified retroviral vectors in embryonal stem cells and embryonal carcinoma cells. J Virol 71, 9466–74 (1997).

19. Gibson, D. G. et al. Enzymatic assembly of DNA molecules up to several hundred kilobases. Nat Methods 6, 343–345 (2009).

20. Ewels, P. A. et al. The nf-core framework for community-curated bioinformatics pipelines. Nat Biotechnol 38, 276–278 (2020).

21. Robinson, M. D., McCarthy, D. J. & Smyth, G. K. edgeR: a Bioconductor package for differential expression analysis of digital gene expression data. Bioinformatics 26, 139–40 (2010).

22. Oh, H. M. et al. STAT3 protein promotes T-cell survival and inhibits interleukin-2 production through up-regulation of class O forkhead transcription factors. Journal of Biological Chemistry 286, 30888–30897 (2011).

23. Chetoui, N., Boisvert, M., Gendron, S. & Aoudjit, F. Interleukin-7 promotes the survival of human CD4+effector/memory T cells by up-regulating Bcl-2 proteins and activating the JAK/STAT signalling pathway. Immunology 130, 418–426 (2010).

24. Tripathi, P. et al. STAT5 Is Critical To Maintain Effector CD8+ T Cell Responses. The Journal of Immunology 185, 2116–2124 (2010).

25. Seo, H. et al. TOX and TOX2 transcription factors cooperate with NR4A transcription factors to impose CD8+ T cell exhaustion. Proc Natl Acad Sci U S A 116, 12410–12415 (2019).

26. Preston, G. C., Feijoo-Carnero, C., Schurch, N., Cowling, V. H. & Cantrell, D. A. The Impact of KLF2 Modulation on the Transcriptional Program and Function of CD8 T Cells. PLoS One 8, 1–16 (2013).

27. Fang, F. et al. Human Transcription Factor KLF3 Maintains T Lymphocyte Quiescent Phenotype Via Inhibiting SHP-1 Expression. Blood 126, 3426–3426 (2015).

28. Gattinoni, L., Speiser, D. E., Lichterfeld, M. & Bonini, C. T memory stem cells in health and disease. Nat Med 23, 18–27 (2017).

29. Chew, G. M. et al. TIGIT Marks Exhausted T Cells, Correlates with Disease Progression, and Serves as a Target for Immune Restoration in HIV and SIV Infection. PLoS Pathog 12, e1005349 (2016).

30. Jackson, Z. et al. Sequential Single-Cell Transcriptional and Protein Marker Profiling Reveals TIGIT as a Marker of CD19 CAR-T Cell Dysfunction in Patients with Non-Hodgkin Lymphoma. Cancer Discov 12, 1886–1903 (2022).

31. Schluns, K. S., Kieper, W. C., Jameson, S. C. & Lefrançois, L. Interleukin-7 mediates the homeostasis of naïve and memory CD8 T cells in vivo. Nat Immunol 1, 426–32 (2000).

